# CLAN: the CrossLinked reads ANalyais tool

**DOI:** 10.1101/233841

**Authors:** Cuncong Zhong, Shaojie Zhang

**Author notes:** Corresponding Author: Email addresses: CZ, SZ.

## Abstract

The crosslinked RNA sequencing technology ligates interacting RNA strands followed by next-generation sequencing. Mapping of the resulting duplex reads allows for functional inference of the corresponding intramolecular/intermolecular RNA-RNA interactions. However, duplex read mapping remains computationally challenging, and the existing best-performing software fails to map a significant portion of the duplex reads. To address this challenge, we develop a novel algorithm for duplex read mapping, called CrossLinked reads ANalysis tool (CLAN). CLAN demonstrates drastically improved sensitivity and high alignment accuracy when applied to real crosslinked RNA sequencing data. CLAN is implemented in GNU C++, and is freely available from http://sourceforge.net/projects/clan-mapping.

## Background

Non-coding RNA (ncRNA) is RNA that do not code for protein; instead, it performs various biological functions such as expression regulation, modification, and catalysis *etc.* [1, 2]. Many of these functions are made possible through the folding into specific RNA structures. For example, the long non-coding RNA (lncRNA) HOTAIR requires a structural basis to expose its PRC2 (polycomb repressive complex 2)-binding motif to properly perform its biological function [3]. Some other ncRNA functions are mediated through RNA-RNA interaction (RRI); e.g. the microRNA (miRNA) interacts with its target mRNA through sequence complementarity and regulates the corresponding gene expression level through regulating the stability of the targeted mRNA [4]. Another important RRI is the binding of the U1 small nuclear RNA (snRNA) and the pre-mRNA, as well as other snRNAs (U2, U4, U5, and U6) recruited during the formation of the spliceosome [5]. As a result, genome-wide study of ncRNA secondary structure and RRI can provide valuable insight to the function of the transcriptome.

Genome-wide RNA secondary structure and RNA-RNA interaction were traditionally studied computationally. The RNA folding [6-8] and co-folding [9, 10] algorithms seek to find the Minimum Free Energy (MFE) structure of a single RNA molecule or RNA duplex, respectively. Sequence-based RRIs, such as miRNA-mRNA interaction, were also computationally predicted using sophisticated computational models [11] that summarize sequence complementarity [12] and site accessibility [13] information. Unfortunately, the existing free energy model [14] and sequence-based interaction model remain insufficient to characterize the complex molecular dynamics, and the computationally predicted RNA secondary structure and RRI remain imprecise. To obtain more accurate RNA secondary structurome, RNA chemical probing technique was coupled with the next-generation sequencing (NGS) technology to allow for genome-wide RNA secondary structure probing; specific technologies include SHAPE-Seq [15], PARS [16], and FragSeq [17] *etc.* However, the NGS-empowered RNA probing technology remains incapable of studying genome-wide RRIs.

Recently, an NGS-based crosslinked RNA sequencing technology was developed to directly probe genome-wide ncRNA secondary structures and RRIs simultaneously (Figure 1). This technology first protects interacting RNA strands (Figure 1, cyan boxes), followed by fragmentation of the RNA molecules. After fragmentation and size selection (or immunoprecipitation if the RNA strands are bond to protein), the protected interacting RNA strands are enriched. The technology then either chemically or radioactively crosslinks the interacting RNA strands, linearizes the product, and adopts standard library preparation and sequencing steps to generate duplex reads (Figure 1). The crosslinked RNA sequencing technology has been applied to different model organisms, and is currently mature enough for human. Specific technologies differ in experimental protocol and biological application, with examples including CLASH [18, 19], iPAR-CLIP [20], MARIO [21], hiCLIP [22], RPL [23], PARIS [24], and LIGR-Seq [25] *etc.* Some of the above methods rely on immunoprecipitation to enrich specific RRIs; for example, CLASH uses the Argonaute (AGO) antibody to specifically pull down interacting miRNA and mRNA [19]. Others are protein-independent and can study transcriptome-wide intramolecular and intermolecular RRIs (e.g. PARIS [24] and LIGR-seq [25]). Mapping of the duplex reads against the reference genome reveals the genomic locations of the two interacting RNA strands (also referred to as RNA *arms*). Intuitively, an intramolecular RRI corresponds to a stem/helix secondary structure in a single RNA molecule, and an intermolecular RRI corresponds to a potential binding site of two interacting RNA molecules.

**Figure 1:**
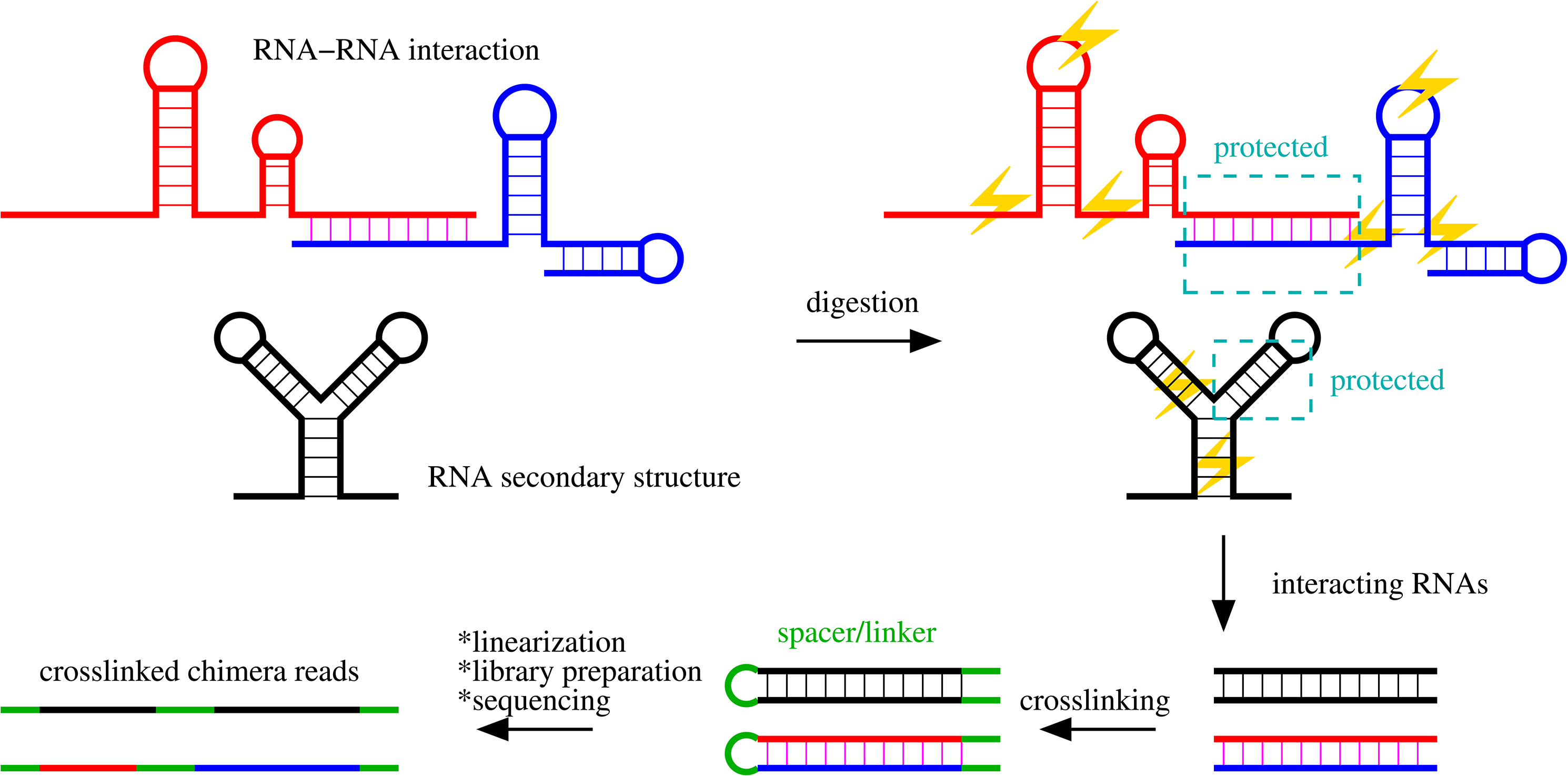
An overview of the duplex read generation process. Interacting RNA strands could be either intermolecular (red and blue RNAs) or intramolecular (black RNA). The interacting RNA strands are either protected by chemical reagents (e.g. psoralen) or ligated to interacting proteins (e.g. AGO or Staufen 1) (cyan boxes). Then, RNase is applied to digest the RNAs and enrich the interacting RNA strands. Consecutively, the enriched RNA strands are crosslinked (with potential incorporation of spacer sequences, internal green segments) and with barcode/adaptors ligated to the ends as linker sequences (flanking green segments). The crosslinked RNA strands are then linearized and subject to standard library preparation and sequencing steps. The resulting duplex reads, in general, have a common architectural pattern of “linker-RNA arm-spacer-RNA arm-linker”.

Unfortunately, the seemingly straightforward analysis strategy is currently hampered by our inability for generating comprehensive and high-quality mapping of the duplex reads. Read mapping is considered as the main cause of information loss, and one of the major challenges in crosslinked RNA sequencing data analysis [26]. To understand the unique computational challenge in duplex read mapping, note the “linker-RNA arm-spacer-RNA arm-linkerˮ pattern of a duplex read (Figure 1). The *linker* sequences (Figure 1, flanking green segments) are often ligated to the termini of the cDNA libraries to assist with PCR amplification, and they can be easily removed with existing trimming software when linker sequence library is given. However, the *spacer* sequence (Figure 1, internal green segment) is more difficult to detect. The mechanism for the inclusion of the spacer sequence has not been thoroughly discussed, and possible reasons could be either biological, experimental, or even artificial (depending on specific experimental protocol). For example, in CLASH [19], the spacer sequence may due to post-transcriptional modification of one of the interaction partners (e.g. oligoadenylation) [27]. While in hiCLIP, the spacer sequence corresponds to an adaptor sequence that is linked to the 5’ end of one of the interaction partners [22]. Moreover, during the crosslinking step, short oligonucleotides may also be crosslinked between the two interacting RNA strands due to opportunistic spatial proximity and become the spacer sequence. In some cases (such as hiCLIP), knowing the adaptor sequence is possible to identify the spacer sequence, however it would inevitably complicate the entire analysis pipeline. In other cases (such as CLASH and proximity-driven crosslinking), it is very difficult to reliably detect the spacer sequence. Currently, existing dedicated analysis pipelines set up a hard cutoff for the length of the spacer sequence (e.g. 4nt for CLASH [27] and 10nt for PARIS [24]). The spacer length cutoff is empirical and highly technology-specific, which may filter out valid duplex reads with longer spacer sequences [27] and poses heavy burden on the users to supply correct parameter. While for general-purpose read-mappers such as BWA [28], BOWTIE2 [29], and STAR [30], most of them can only automatically truncate end sequences (also known as soft-clipping), but not internal sequences [31]. Or, as in STAR [30], it requires the user to supply a hard cutoff to soft-clip internal sequence, which is difficult to estimate and could lead to low sensitivity for the above mentioned reason. The consequence of improper handling of a spacer sequence is that the spacer sequences will be treated as excessive sequencing errors, which makes the mappers discard the corresponding duplex read, and subsequently leading to low mapping sensitivity.

In addition to the spacer sequence, two other reasons also make duplex-read mapping challenging. First, many alignment/mapping tools require long seed match to initialize the alignment to ensure high computational efficiency. For example, the current version of BLASTN requires a pair of non-overlapping 11-nt gapped seeds to initiate an alignment [32]. Such a requirement implies that each RNA arm needs to be at least 22nt long, which disqualifies many (partial) miRNA arms. The seed-length requirement may be more stringent in mapping tools (e.g. BOWTIE2, which requires 28nt seed-matches by default). Second, the layout of a crosslinked RNA sequencing read can be different from regular RNA-seq reads. The majority of RNA-seq mapper are restricted to detecting splicing events, where the two exonic sequences must be mapped to the same chromosome. However, in crosslinked RNA sequencing data, inter-chromosomal RNA arms could be derived from inter-molecular RRIs, making the read difficult to be handled by programs restricted to detecting splicing events.

Unfortunately, all issues mentioned above have not been fully recognized, and most existing studies still adopt general-purpose mapper such as BLASTN [32], BOWTIE2 [29], or STAR [30], leading to extremely low read-mapping rate (e.g. 3% as reported in CLASH [19]). Other dedicated analysis pipelines, such as Hyb [27] designed for CLASH datasets and Aligator [25] for LIGR-Seq datasets, also call the existing alignment/mapping tools as its read-mapping subroutine and are therefore subject to the abovementioned issues. As a result, to the best of our knowledge, no existing alignment/mapping software suitable for duplex read mapping is available on the market.

We attempt to address the existing limitations in crosslinked RNA sequencing read mapping by formulating a novel read-mapping problem. We respect the fact that each duplex read may contain (random or unidentified) linker/spacer sequences; and only partial sequence of the read is informative and corresponds to interacting RNA strands. With this intuition, we seek to identify two non-overlapping substrings of the read, where each substring can be mapped to the reference genome with less than a given number of edits (sequencing errors/polymorphisms), and that the total length of the two substrings is maximized. With this formulation, no information regarding the adaptor sequence nor spacer length cutoff is expected from the user, and both spacer and linker sequences will be detected automatically. We construct Burrows-Wheeler Transformation (BWT) and the corresponding FM-index on the reference genome, and perform exhaustive search of all prefixes of the read to find potential mappings. We then merge and chain (using dynamic programming) the resulting mapped substrings, to identify the nonoverlapping pair of substrings with the maximized total length. Because our algorithm respects the existence of the linker/spacer sequence and performs exhaustive search of all mappings, we anticipate its much higher mapping sensitivity. Also, with heuristics (detailed in the Methods section), our algorithm is expected to perform efficiently in practice.

We implemented the algorithm into software called CLAN (the <u>C</u>ross<u>L</u>inked reads <u>AN</u>alysis tool). We benchmarked CLAN with popular alignment/mapping tools BLASTN [32] and STAR [30]; we selected BLASTN as the representative of alignment tools for its popularity, and STAR as the representative of mapping tool both for its high mapping performance [31] and its ability to map chimeric reads. We benchmarked the three programs on four different crosslinked RNA sequencing datasets including CLASH [19], hiCLIP [22], PARIS [24], and LIGR-Seq [25], for all of them were derived from human samples. We found that CLAN was capable of mapping much more reads than BLASTN and STAR, with the most significant improvement being observed from the CLASH dataset, where >90% of the reads were uniquely mapped by CLAN. Furthermore, the mapping locations predicted by CLAN are also highly accurate, with >90% of them being consistent with those predicted by BLASTN. Compared to BLASTN and STAR, CLAN requires ∼30% more physical memory; however, the requirement (~37G for human genome) can be easily accommodated by current computing facility. CLAN runs hundreds of times faster than BLASTN, and only 2-3X slower than STAR; the extra running time can be easily accommodated by introducing extra computing units. In summary, CLAN is a powerful tool for crosslinked RNA sequencing read mapping with high sensitivity, high accuracy, and high speed. CLAN is implemented in GNU C++, and is freely available from http://sourceforge.net/projects/clan-mapping.

## Results

We selected four crosslinked RNA sequencing datasets that were generated by different groups and different technologies (CLASH [19], hiCLIP [22], PARIS [24], and LIGR-Seq [25]) to benchmark the performance of CLAN together with BLAST [32] and STAR [30]. We only analyzed the mappings of the first 2.5 million reads from each dataset, as BLAST was unable to finish the mapping of the entire datasets within a reasonable time; the in-total 10 million reads dataset is sufficiently large to generate statistically meaningful conclusions for a benchmark purpose. Each dataset was quality-trimmed using Trimmomatic [33] under default parameters. Details on the benchmark dataset are included in Table 1.

**Table 1:** Summary of the benchmark datasets

| Technology | SRA Accession | Ave. Read Len | Num. Reads |
| --- | --- | --- | --- |
| **CLASH** | SRR959751 | 55 | 2,313,448 |
| **hiCLIP** | ERR605257 | 196 | 2,331,483 |
| **PARIS** | SRR3194440 | 100 | 2,476,786 |
| **LIGR-Seq** | SRR3361013 | 102 | 2,223,699 |

For each dataset, the first 2.5 million reads were initially selected for the analysis. Column “Num. Reads” corresponds to the number of reads survived after quality trimming.

The reads were mapped against the human reference genome (version hg38) using CLAN, BLASTN, and STAR. CLAN was run under default parameters (see the Methods section for details). To make BLASTN (version 2.6.0+) run within a reasonable time, the maximum number of target sequence to search was set to 20 (option ‘-max_target_seqs 20’); other default parameters were used. For STAR (version 2.5.3a), we also set the maximum number of mappings to report as 20 (option ‘--outFilterMultimapNmax 20’); we also turned on STAR’S chimera read mapping mode (option ‘--chimSegmentMin 5’, which requires at least 5nt for each reported RNA arm). Other default parameters were used for STAR. All programs were run with 8 threads. All experiments were performed on an in-house server equipped with Intel(R) Xeon(R) CPU E7-4850 v4 @ 2.10GHz and 1T RAM.

### Computational efficiency of CLAN

We summarize the wall-clock running time and peak memory consumption of CLAN, BLASTN, and STAR in Table 2. STAR demonstrated the highest speed among the three programs tested. CLAN was ~2-3X slower than the read mapping tool STAR, because CLAN exhaustively searched all possible matchings. On the other hand, BLASTN required a much longer running time (~300X slower in the worst case) than CLAN and STAR, which makes BLASTN inappropriate for routine analysis of large sequencing datasets. By extrapolation, BLASTN may require ~42 days to map the entire LIGR-Seq dataset (~50 million reads) with 8 CPUs; while CLAN would take ~3hrs for the same task. The memory consumption of the three software was similar in the worst-case scenario, with CLAN required ~30% more memory than the other software.

**Table 2:** Wall-clock running time and peak memory consumption of CLAN, BLASTN, and STAR

| Technology | CLAN |  | BLASTN |  | STAR |  |
| --- | --- | --- | --- | --- | --- | --- |
|  | Time | RAM | Time | RAM | Time | RAM |
| **CLASH** | 6m45s | 37G | 9m1s | 4G | 3m58s | 29G |
| **hiCLIP** | 11m36s | 37G | 20h48m26s | 11G | 6m13s | 29G |
| **PARIS** | 11m22s | 37G | 7h56m9s | 26G | 5m20s | 29G |
| **LIGR-Seq** | 9m17s | 37G | 43h23m44s | 13G | 3m35s | 29G |

All programs listed above were run with 8 threads.

### CLAN mapped more duplex reads than BLASTN and STAR

We summarize the read mapping results of CLAN, BLASTN, and STAR on CLASH, hiCLIP, PARIS, and LIGR-Seq datasets in Figure 2. In all four cases, CLAN mapped more reads than BLASTN and STAR. CLAN’s high sensitivity is most apparent for the CLASH dataset. Specifically, the novel rate of CLAN mapping (defined by the number of unique CLAN mapping over the total number of mapped reads) was 94.9% (CLASH), 24.7% (hiCLIP), 30.5% (PARIS), and 15.8% (LIGR-Seq). The majority of the reads that can be mapped by BLASTN or STAR can also be mapped by CLAN, with CLAN missing only 3 CLASH reads, 3 hiCLIP reads, and 50 LIGR-Seq reads that can be mapped by BLASTN; CLAN also mapped all reads that can be mapped by STAR. Surprisingly, the mapping of BLASTN and STAR was not entirely consistent, with the Jaccard similarity coefficient being 0.157 (CLASH), 0.249 (hiCLIP), 0.440 (PARIS), and 0.761 (LIGR-Seq). Although STAR was significantly faster than BLASTN, it mapped 51.4% less CLASH reads, 65.7% less hiCLIP reads, 53.2% less PARIS reads, and 16.0% less LIGR-Seq reads than BLASTN, suggesting that the sensitivity of the mapping tool may be compromised for its higher speed. Overall, the reads mapped by CLAN was almost a superset of the union of the BLASTN and STAR mappings, thus CLAN demonstrated the highest mapping sensitivity and robustness among all three programs tested.

**Figure 2:**
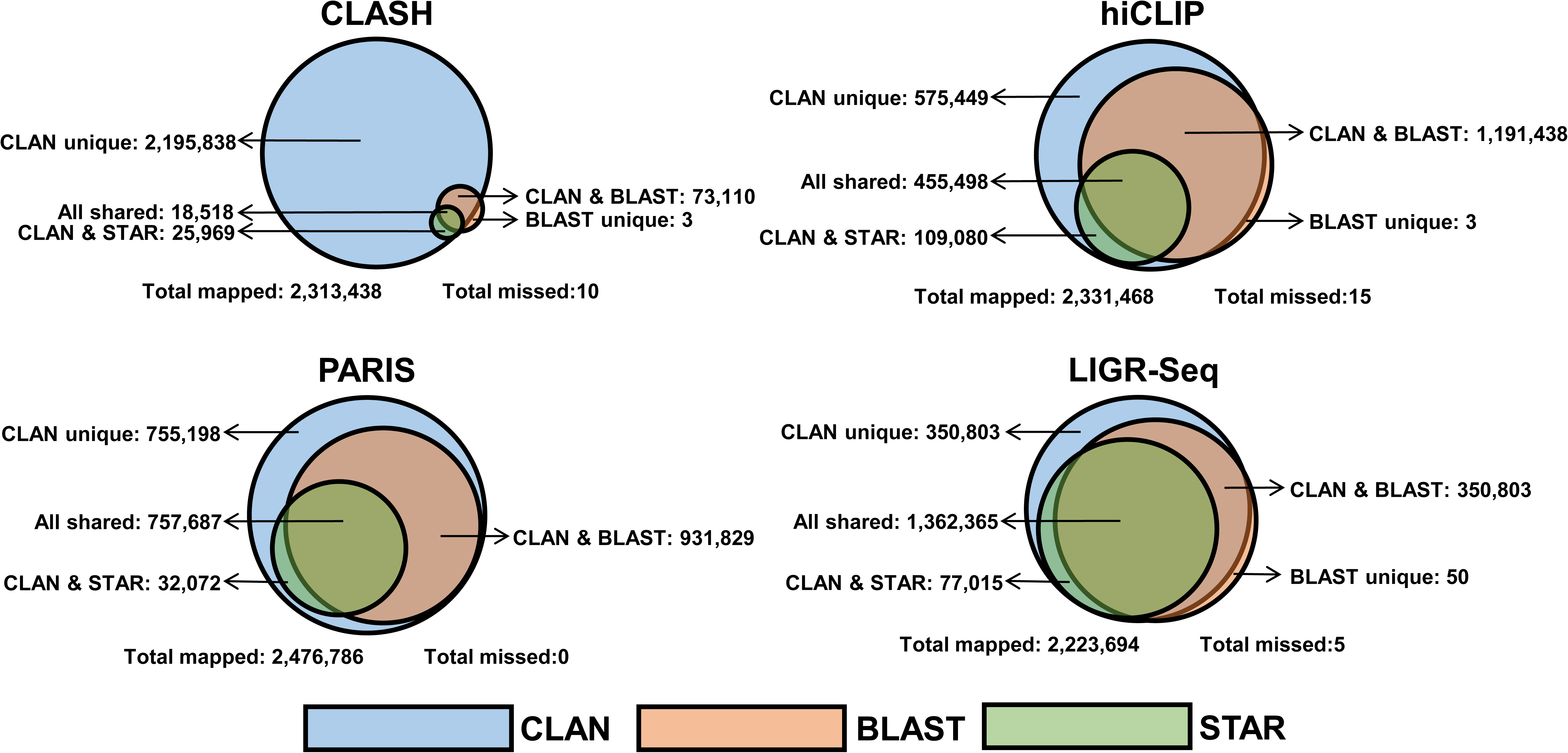
Venn diagram of the read-mapping results generated by CLAN (blue), BLAST (red), and STAR (green) on the CLASH, hiCLIP, PARIS, and LIGR-Seq datasets. “CLAN unique” indicates reads that are uniquely mapped by CLAN and not by BLASTN nor STAR; similarly for “BLAST unique”. No read is uniquely mapped by STAR. “CLAN & BLAST” indicates reads mapped by both CLAN and BLASTN but not by STAR; similarly for “CLAN & STAR”. No read is mapped by both BLASTN and STAR but not by CLAN. “All shared” indicates reads that are mapped by all three programs. “Total mapped” indicates the reads that can be mapped by at least one of the programs, and “Total missed” indicates the reads that cannot be mapped by any of the programs.

### CLAN accurately mapped duplex reads

To analyze the mapping accuracy of CLAN, we compare the mapping locations generated by CLAN, BLASTN, and STAR. We decompose the CLAN mappings into four categories by comparing the mapping locations with those predicted by BLASTN (or STAR). For each CLAN-mapped read, if the mapping location was identical to those predicted by BLASTN (or STAR), we consider the mapping as *consistent*. Or, if the mapping location overlapped with those predicted by BLASTN (or STAR) for >60% of its total length, we consider the mapping as *overlap*. Third, if the mapping location overlapped with those predicted by BLASTN (or STAR) for ≤60%, we consider the mapping as *inconsistent*. Finally, if the mapping was uniquely generated by CLAN, we consider the mapping as *novel*. Define *concordance rate* by using the following formula:

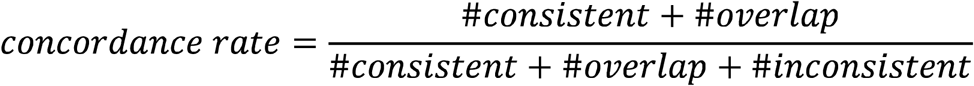

The mapping location comparison between CLAN and BLASTN is summarized in Figure 3. Overall, the CLAN prediction was highly consistent with BLASTN prediction, with concordance rates being 98.0%, 95.2%, 94.4%, and 92.3% for the CLASH, hiCLIP, PARIS, and LIGR-Seq datasets, respectively. We perform the same comparison between CLAN and STAR predictions and the result is summarized in Figure 4. The concordance rates were also high for all four datasets; specifically, 87.7%, 85.8%, 79.2%, and 81.3% for the CLASH, hiCLIP, PARIS, and LIGR-Seq datasets, respectively. Overall, both comparisons suggest that the mapping locations predicted by CLAN were concordant with the existing software, which further indicates that CLAN correctly maps the duplex reads. Furthermore, note that the concordance rates between CLAN and BLAST, for all four datasets, were higher than those between CLAN and STAR. Since BLASTN adopts the traditional alignment algorithm for its seed-and-extend strategy, the mapping generated by BLASTN is, in most cases, considered as the most accurate one among those generated by the existing mapping/alignment software. The results thus suggest that CLAN was capable of generating more accurate mapping than STAR. In summary, the mapping predicted by CLAN is highly concordant to that predicted by BLASTN, and is more accurate than that predicted by STAR.

**Figure 3:**
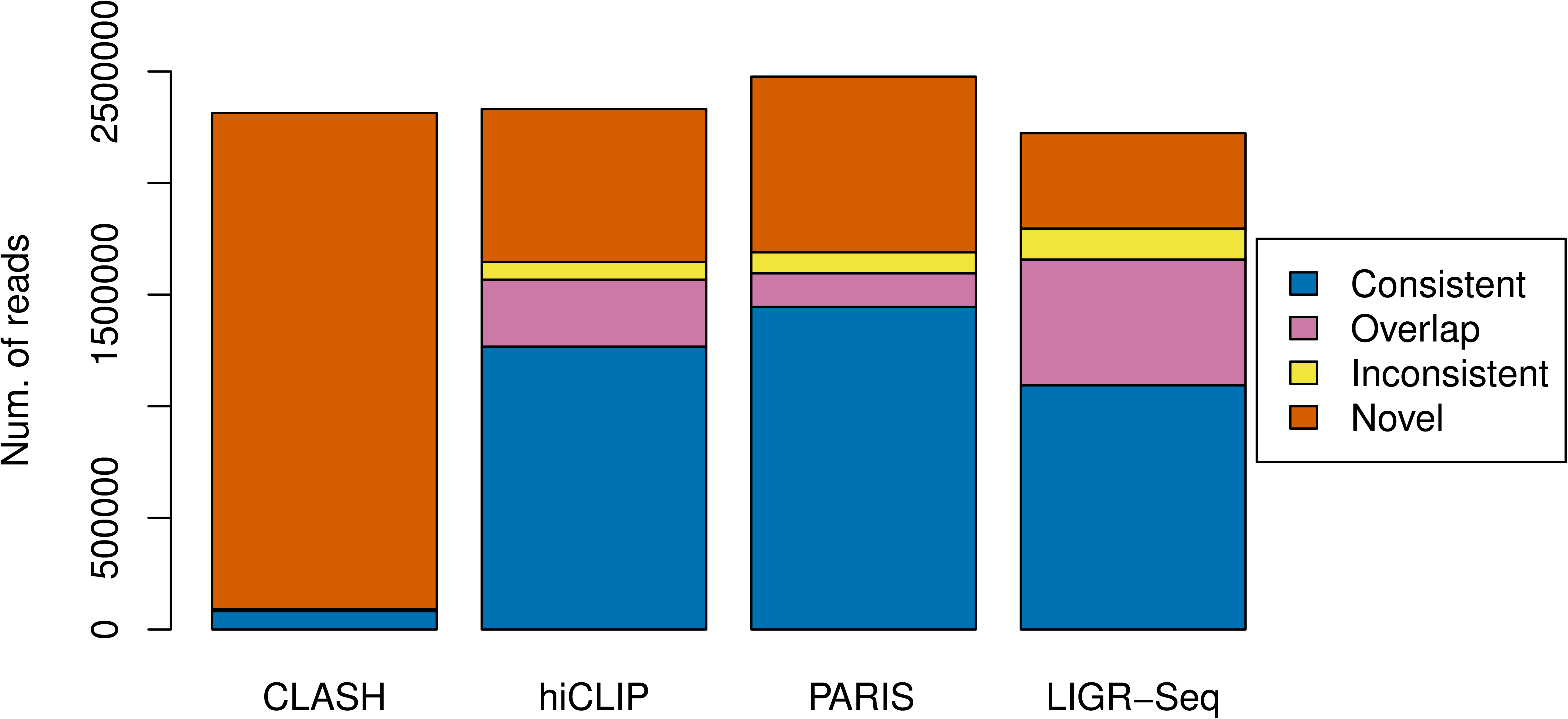
Decomposition of the CLAN read mapping for the CLASH, hiCLIP, PARIS, and LIGR-Seq datasets by comparing with BLASTN mappings. “Consistent”: reads whose mapping locations predicted by CLAN and BLASTN are identical. “Overlap”: reads whose mapping locations predicted by CLAN and BLASTN overlap for >60% of the total length. “Inconsistent:” reads mapped by both CLAN and BLASTN but do not overlap or overlap for ≤60% of the total length. “Novel”: reads uniquely mapped by CLAN.

**Figure 4:**
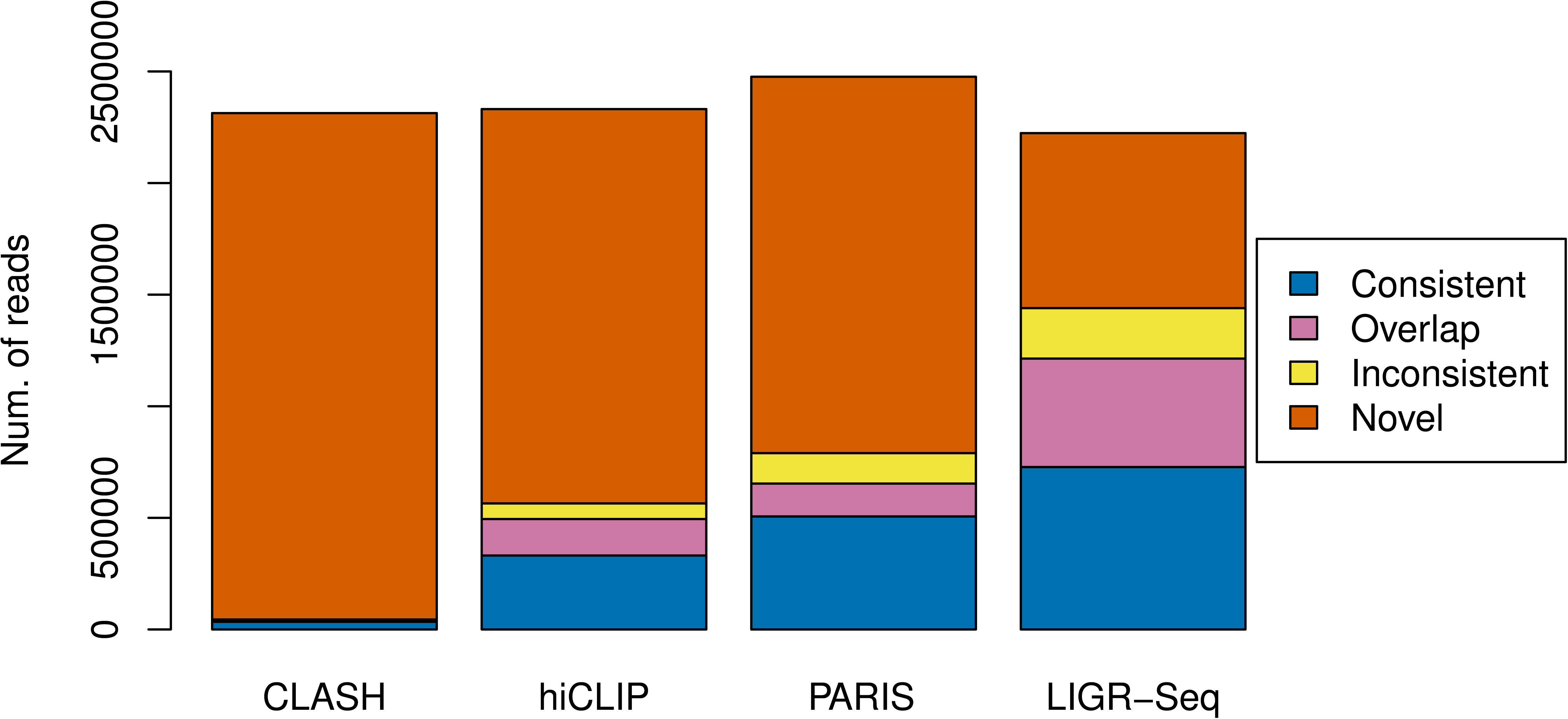
Decomposition of the CLAN read mapping for the CLASH, hiCLIP, PARIS, and LIGR-Seq datasets by comparing with STAR mappings. “Consistent”: reads whose mapping locations predicted by CLAN and STAR are identical. “Overlap”: reads whose mapping locations predicted by CLAN and STAR overlap for >60% of the total length. “Inconsistent reads mapped by both CLAN and BLASTN but do not overlap or overlap for ≤60% of the total length. “Novel”: reads uniquely mapped by CLAN.

### Mechanisms for CLAN to generate overlapping and inconsistent mappings

We then take a deeper look at the cases where CLAN generated overlapping and/or inconsistent mappings when compared to BLASTN predictions. We summarize the major mechanisms in Figure 5. Figure 5(A) demonstrates the major mechanism for CLAN to generate overlapping mapping. In BLASTN, the read was aligned to the reference with two mismatches (Figure 5, red “C” and “A”). Since CLAN only allowed for one error/polymorphism (by default, see the Methods section for more details), the alignment of the short sequence (“CGG”, blue) after the second mismatch was discarded, which made the mapped region shorter than BLASTN’s mapping.

**Figure 5:**
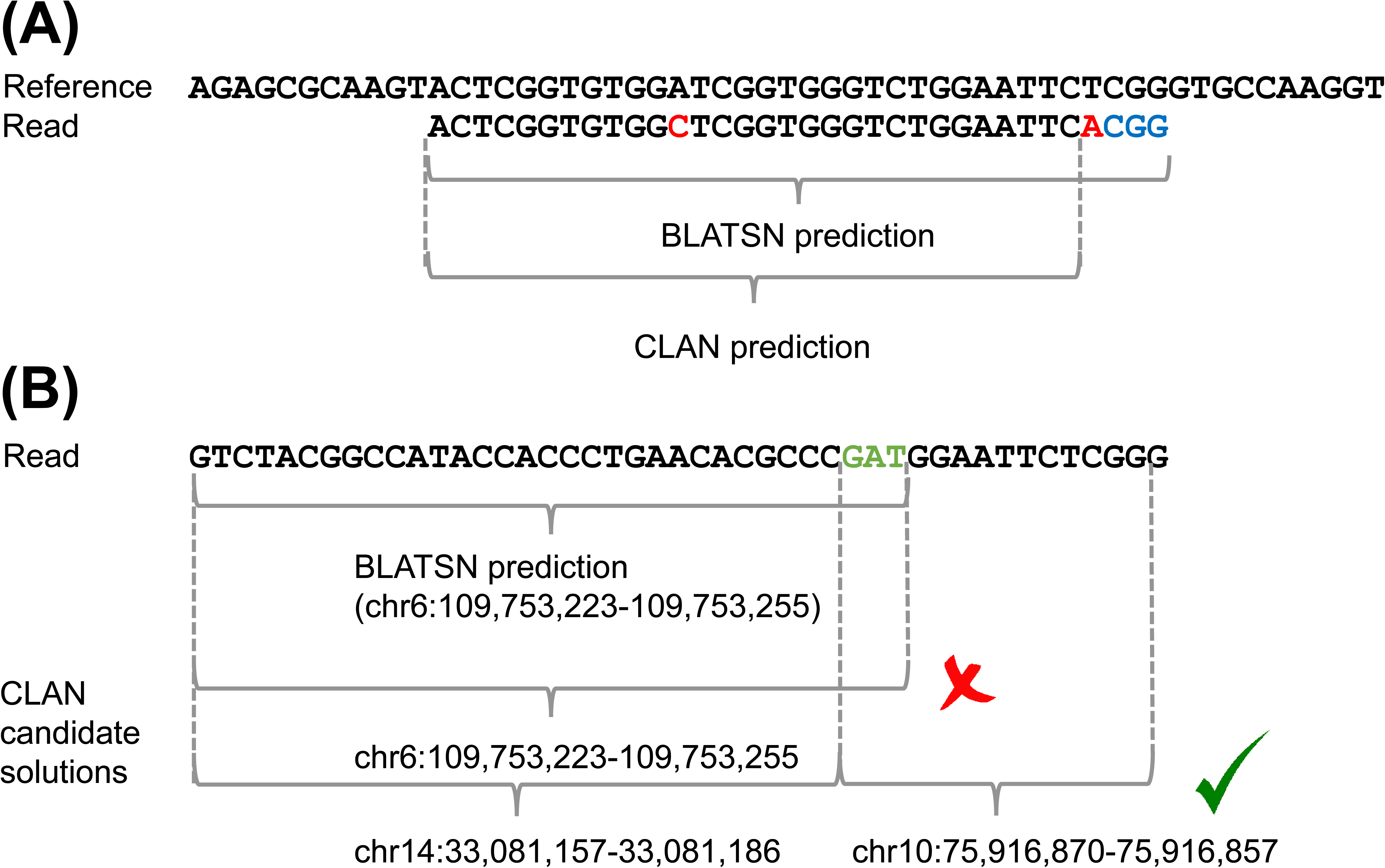
Mechanisms for CLAN to generate “Overlap” or “Inconsistent” mappings (as compared to BLASTN). (A) The major reason for “Overlap” mappings. Since CLAN only allows a given number of errors in the mapped region, additional alignment segments that lead to more errors than allowed will be discarded. In this example where only one error is allowed, CLAN has already detected one error (the red “C” in read), and therefore it discards the additional alignment segment (blue segment in read, which can be detected by BLASTN) since the inclusion of such a segment leads to an additional error (the red “A” in read). (B) The major reason for “Inconsistent” mappings. CLAN identifies the mapping of the prefix (up to the green segment “GAT”, inclusive) against chr6 (the gray brace in the second row), and therefore discards the mapping of the subsequent prefix (up to the green segment “GAT”, exclusive) to the same genomic positions (chr6) as redundancy. At this point, the substring prefix only maps to chr14 (the first gray brace in the third row). During chaining, the read is optimally decomposed into two RNA arms (indicated by the last two gray braces in the third row) and the mapping of the prefix (up to the green segment “GAT”, exclusive) is thus directed to chr14. On the other hand, BLASTN only identifies the mapping of the longer prefix (green segment “GAT”, inclusive) against chr6 (gray brace in the first row).

Figure 5(B) illustrates the major mechanism for CLAN to generate inconsistent mapping. In this case, the read can be ambiguously treated as a single-arm read (Figure 5(B), single gray brace) or a double-arm read (Figure 5(B), double gray braces); when treated as a double-arm read, some common sequences (Figure 5(B), green sequence “**GAT**”) were shared at the flanking regions of the two arms. BLASTN treated the read as a single-arm read by assigning the shared sequence to the first arm, while CLAN treats the read as a double-arm read by assigning the shared sequence to the second arm. Since when performing the backward exhaustive BWT search, CLAN first identified the mapping of the first-arm prefix (up to the shared sequences “**GAT**”, inclusive) to chr6. In consecutive BWT searches, CLAN considered the mapping of the shorter prefix (up to the shared sequence “**GAT**”, exclusively) to the same location at chr6 as redundant, and subsequently discarded such a mapping location. An alternative mapping location (chr14) remained for the shorter prefix. As a result, after chaining, CLAN reported the read as a double-arm read, with both arms respectively mapped to chr14 and chr10, completely different from the mapping to chr6 as reported by BLASTN. The mapping location to chr14 predicted by CLAN also appeared in BLATSN’s high-scoring list (8^th^ place, 100% identity), as well as in the corresponding BLAT [34] search (2^nd^ place, 100% identity). As a result, the inconsistency of mapping was primarily due to different programs’ preferences in assigning read mapping, but not an error of CLAN. We also note that one can avoid such a mapping inconsistency by setting higher duplex mapping cost (see more details in the Methods section), such that CLAN will favor more in treating the read as single-arm read.

### CLAN analysis of the CLASH data revealed potential novel miRNA-mRNA interactions

Here, we showcase biological applications of CLAN in discovering novel miRNA-mRNA interactions by analyzing the entire CLASH dataset SRR959751, which was generated from the Flp-In T-REx 293 cell line derived from human kidney stem cells [19]. The entire dataset contains 48,695,407 reads in total after Trimmomatic [33] trimming (using the same criterion as described above). CLASH mapped 48,681,395 (99.97%) of them. The mapped reads were further annotated using ANNOVAR (version 2017-07-17; under the gene annotation mode) [35] with RefSeq hg38 as the genome annotation database. As recommended by the CLASH authors [27], the annotation was made strand-specific and only the mappings on the annotated transcript strand was considered. Among all mapped reads, 11,993,182 (24.63%) of them have one of both arms mapped to an annotated microRNA; furthermore, 11,819 reads have one of their both arms mapped to an annotated microRNA and at the same time the other arm mapped to an annotated 3’UTR of a gene. These reads were clustered based on their mapped locations, and finally revealed 1,042 unique miRNA-mRNA interactions (see Supplementary Table S1).

As an example, we performed further analysis on interactions relating to miR-10a. In total, 46 miR-10a-mRNA interactions were supported by at least one CLASH duplex read. We used RNAcofold [10] to perform RNA dimer binding analysis on these predicted interactions (as in ViennaRNA Package [6] v2.4.3, with parameter ‘-a’ to compute heterodimer energy). The mature miRNA sequences were taken from miRBase [36]. For the mRNA sequences, since CLASH is unable to reveal single-nucleotide-resolution information regarding the interacting RNA arms, the mRNA sequences were extended by 10nt towards both the 5’ and 3’ ends (as recommended in Hyb [27]). Among the 46 interactions, only 5 (10.8%) were predicted by the original CLASH analysis (using BLASTN as the mapping tool), and only 3 (6.5%) were predicted by TargetScan [12]. The 46 interactions were sorted based on the number of supporting duplex reads (Supplementary Table S2). The 15 interactions with more than 5 supporting reads are summarized in Table 3.

**Table 3:** The predicted mRNA targets of miR-10a with >5 supporting duplex reads identified by CLAN re-analysis of the CLASH (SRR959751) dataset

| Target | Chrom | Start | End | Strand | #Reads | dG | CLASH | TargetScan |
| --- | --- | --- | --- | --- | --- | --- | --- | --- |
| **RWDD2A** | chr6 | 83197331 | 83197369 | + | 521 | -8.57 | N | N |
| **TNS1** | chr2 | 217803240 | 217803277 | - | 385 | -10.99 | N | N |
| **FAM126A** | chr7 | 22942310 | 22942345 | - | 119 | -5.6 | N | N |
| **ASXL2** | chr2 | 25741069 | 25741102 | - | 112 | -5.38 | N | N |
| **RPRD1A** | chr18 | 35992097 | 35992142 | - | 89 | -15.28 | Y | Y |
| **NEBL** | chr10 | 20784131 | 20784165 | - | 88 | -10.49 | N | N |
| **CMYA5** | chr5 | 79800208 | 79800241 | + | 50 | -5.22 | N | N |
| **TRPM7** | chr15 | 50557926 | 50557968 | - | 43 | -6.82 | Y | N |
| **UCP3** | chr11 | 74000325 | 74000358 | - | 36 | -12.52 | N | N |
| **GATAD2B** | chr1 | 153805110 | 153805144 | - | 36 | -12.69 | N | N |
| **DCUN1D5** | chr11 | 103061765 | 103061800 | - | 14 | -5.37 | N | N |
| **HLCS** | chr21 | 36751535 | 36751569 | - | 13 | -8.03 | N | N |
| **WAC** | chr10 | 28621309 | 28621343 | + | 11 | -9.17 | N | N |
| **BRWD3** | chrX | 80673236 | 80673271 | - | 9 | -5.59 | N | N |
| **CTPS1** | chr1 | 41012149 | 41012185 | + | 6 | -6.38 | N | N |

Annotation of fields: “#Reads”: the number of duplex reads supporting the corresponding miRNA-mRNA interaction; “dG”: heterodimer free energy (kcal/mol) predicted by RNAcofold; “CLASH”: whether the interaction was predicted by the original CLASH analysis using BLASTN as the mapper; “TargetScan”: whether the interaction is included in the TargetScan database.

We visualize the corresponding base pairs of these 15 predicted miR-10a-mRNA interactions in Figure 6. As the figure shows, most of the predicted miR-10a-mRNA interactions are facilitated by a large number of inter-arm base pairings. The strongest miR-10a-mRNA binding was observed at the 3’UTR of *RPRD1A,* which corresponds to a free energy of -15.25 Kcal/mol and is supported by 89 CLASH duplex reads. Correspondingly, the miR-10a-RPRD1A interaction is the only one that was both predicted by TargetScan and the previous CLASH analysis. The base pairs formed between miR-10a and *RPRD1A* 3’UTR are also consistent with the existing annotation of the miR-10a seed region. The previous CLASH analysis also predicted the miR-10a-*TRPM7* interaction. The majority of the miR-10a-related interactions listed were not identified by neither TargetScan nor the previous CLASH analysis; and many interactions such as those relating to *TNS1, NEBL, UCP3,* and *GATAD2B* have low free energy and a significantly amount of supporting duplex reads (see Table 3). Surprisingly, the predicted *GATAD2B* 3’UTR target even contains the conserved binding motif (“ACAGGGUA”) of miR-10a, as revealed by the multiple sequence alignment generated from 18 vertebrates (see Supplementary Figure S1). The base pairs formed within the predicted miR-10a-GATAD2B interaction are also consistent with the annotated miR-10a seed region. The other predicted interactions involve base pairs overlapping with the annotated seed region, however the interaction patterns appear to be non-canonical. For example, the binding between miR-10a and *FAM126A* is mediated by 9 consecutive base pairs (with a bulge loop created by a single-nucleotide insert at the *FAM126A* 3’UTR), while only one base pair overlaps with the annotated miR-10a seed region. In summary, these results suggest that current computational methods for miRNA target prediction remain imperfect and may miss many true targets, and coupling experimental data with high-performance analysis tools such as CLAN will reveal a more comprehensive picture of the miRNA-mRNA interactome.

**Figure 6:**
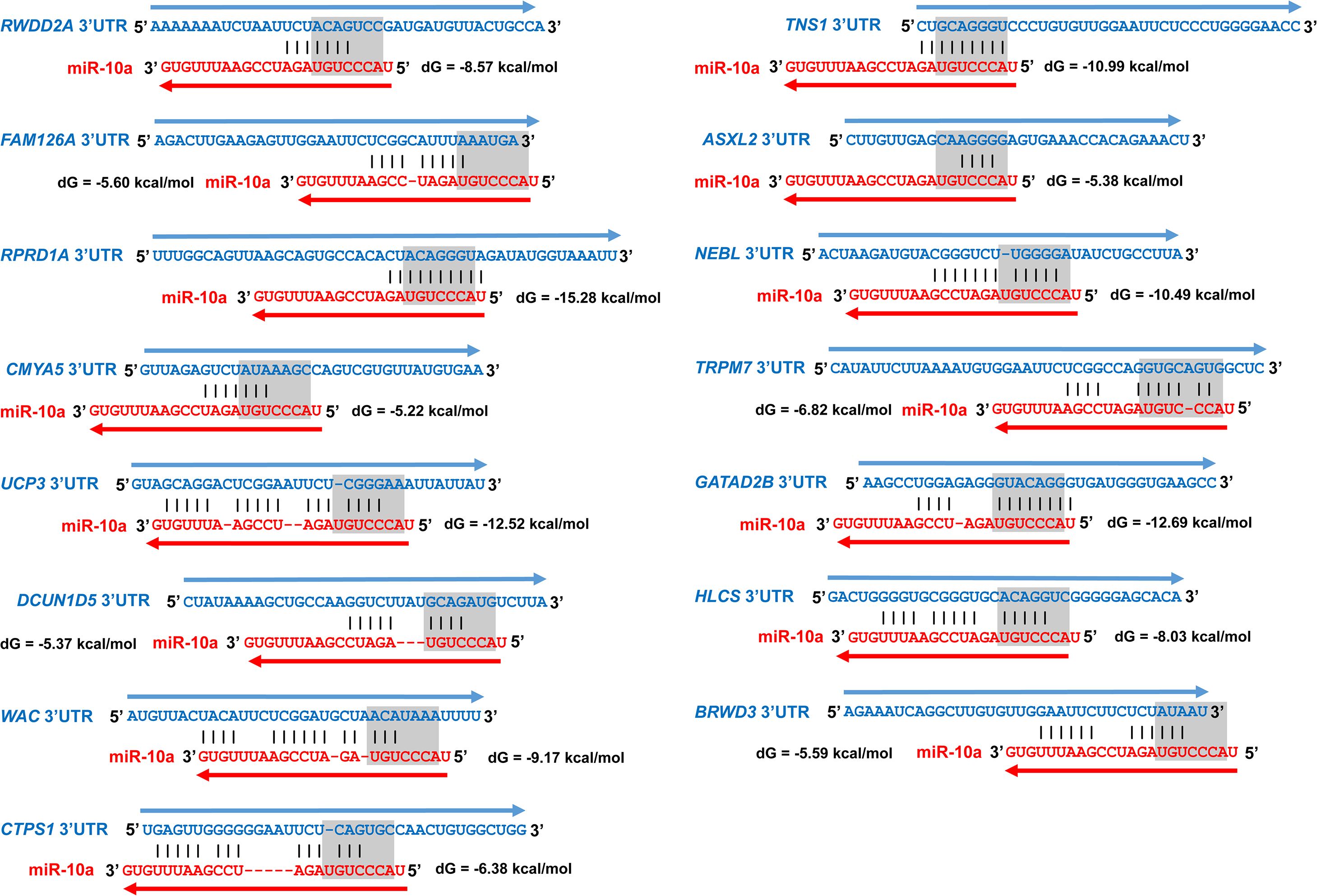
Visualization of miR-10a-mRNA interactions that have at least 5 CLASH duplex read supports. The interactions were predicted using RNAcofold. The red sequences correspond to the mature miR-10a sequence, and the blue sequence corresponds to the extended mRNA target revealed by the CLASH data. The canonical seed region of miR-10a is highlighted by the gray box.

We further analyzed the related pathways associated with the predicted targets of miR-10a to provide additional insights for understanding the biological function of miR-10a. We used Cytoscape (version 3.5.1) [37] to perform gene set enrichment analysis among the 46 predicted miR-10a target genes and their linker genes; the identified interactions and the 10 most significantly enriched pathways are shown in Figure 7 (a complete list of enriched pathways with FDR < 0.01 is available from Supplementary Table S3). In the network shown in Figure 7(A), the blue round nodes correspond to predicted miR-10a targets, the red round nodes correspond to the linker genes among the targets, and the red diamond nodes correspond to hubs (≥10 interactions) of the pathway. The hubs suggest two central biological functions of the network, i.e. ubiquitination (involving hubs *UBB* and *UBC*) and the signal transduction by binding to phosphoserine-containing proteins (involving hubs *YWHAB, YWHAG,* and node *YWHAZ*). The most significantly enriched biological pathways shown in Figure 7(B), i.e. the Hippo signaling pathway and the MAPK signaling pathway, shed lights for our understanding of the cooperation of the two central biological functions in the network. Ubiquitination has been reported as a key regulator of the MAPK signaling pathway [38], and thus can indirectly regulate the Hippo signaling pathway [39]. Interestingly, recent research also revealed that under hypoxia condition, the Hippo signaling pathway can be directly regulated through the *SIAH2* ubiquitin E3 ligase [40]. The hippo signaling pathway play critical roles in restraining cell proliferation and promoting apoptosis, and is critical in stem cell and tissue specific progenitor cell self-renewal and expansion [41]. Its overrepresentation is consistent with the cell line, i.e. the human kidney stem cell, from which the CLASH dataset SRR959751 was generated. The pathway analysis and results shown in Figure 7 suggest miR-10a’s role in regulating cell apoptosis under overgrowth condition, which is also consistent with the aberrant expression of miR-10a in many cancerous cells (e.g. glioblastoma [42], hepatocellular carcinomas [43], colon cancer [44], melanoma [45], breast cancer [45], chronic myeloid leukemia [46], and acute myeloid leukemia [47, 48]). More importantly, more and more evidences have been accumulated to support mir-10a’s direct regulation of the apoptosis pathways under these cancerous conditions [49, 50]. Using CLAN, we were able to identify much more direct targets of miR-10a (40/46 are novel) and construct the complete miR-10a related pathway, which provides much more comprehensive and detailed information for the elucidation of miR-10a’s biological functions.

**Figure 7:**
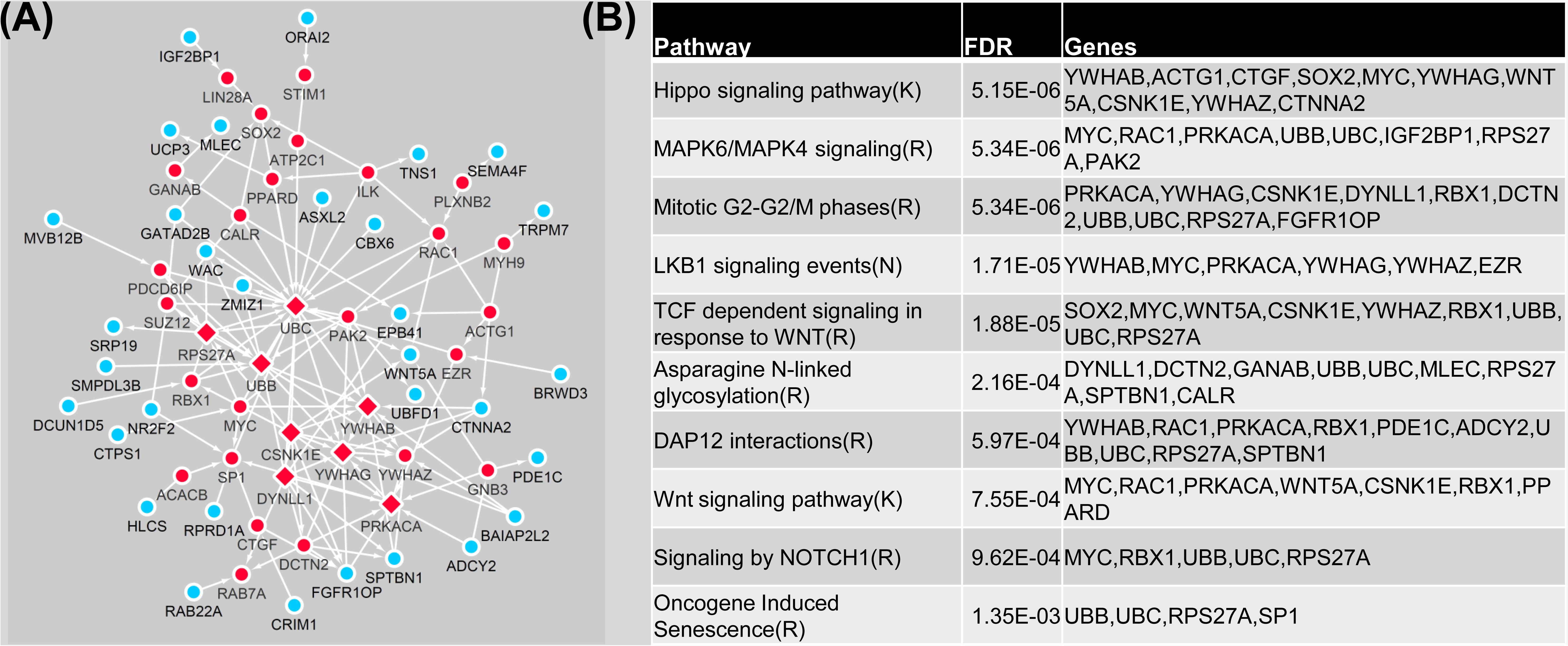
The interaction network and enriched biological pathways identified from the CLAN-predicted miR-10a target genes. (A) The interaction network. Blue round dots correspond to the predicted miR-10a target genes, red round dots correspond to linker genes, and the red diamond dots correspond to hubs which involve at least 10 interactions within the network. (B) The enriched biological pathways (labels in the first column: (K): KEGG, (R): Reactome, and (N): NetPath), the corresponding false discovery rate (FDR), and the involved genes.

## Discussion

In this article, we present a novel algorithm CLAN for duplex read mapping. CLAN was applied to four different datasets generated by CLASH, hiCLIP, PARIS, and LIGR-Seq technologies. Compared to BLASTN, CLAN mapped 96.0%, 29.4%, 31.8%, and 19.2% more reads for the CLASH, hiCLIP, PARIS, and LIGR-Seq datasets, respectively. The same trend was also observed for the comparison between CLAN and STAR. Apparently, the highest improvement over mapping rate by CLAN was observed when analyzing CLASH dataset, because the average read length for the CLAN dataset is the shortest (55bp, see Table 1). For the other datasets that have longer average read lengths, CLAN can still find mapping for more reads, however the improvement was less significant compared to CLASH data. We argue that CLAN remain highly useful even when current sequencing technologies routinely generate longer reads. First, many RNA-RNA interaction intrinsically involving short RNA strands (e.g. miRNA-mRNA interactions studied by CLASH), and the resulting duplex cDNA libraries will inevitably contain the corresponding short RNA arms (i.e. the miRNA that is ~22-28bp long). The analysis of these short RNA arms cannot be made easier with longer reads, and still requires CLAN’s ultra-high sensitivity and accuracy. Second, when the sequencing length grows longer, the running time of BLASTN becomes much longer than CLAN (300X slower, see Table 2); on the other hand, mapping tools such as STAR are fast but cannot match the mapping sensitivity and accuracy of BLASTN. CLAN remains the only available choice with a speed comparable to mapping tools like STAR and a mapping accuracy comparable to alignment tools like BLASTN. As a result, CLAN is a unique and highly useful tool for duplex reads mapping.

We performed deeper analysis on the CLAN mapping of the CLASH dataset and identified potential biological discoveries to showcase CLAN’s real-world applications. Because the limit of the space, we have not present the corresponding findings made when analyzing the hiCLIP, PARIS, and LIGR-Seq datasets. However, both intermolecular and intramolecular RNA-RNA interactions were observed from the reanalysis of PARIS and LIGR-Seq, which could be used in RRI and RNA secondary structure predictions. We note, although CLAN can accurately map duplex reads, it remains incapable of telling whether the mapped duplex reads correspond to real RNA-RNA interactions, as the duplex may be resulted from opportunistic spatial proximity [19]. Experiments or other auxiliary information may be required to confirm the biological relevance of the mappings produced by CLAN.

Currently, CLAN reports all mappings that are equivalently optimal. Because each RNA arm is usually short, multiple genomic locations may be contained in the output. The rationale for this setting is to provide the most comprehensive mapping information to the users of CLAN; and one can devise a tailored strategy to post-process the multi-mapping according to specific research purposes. For example, only may prioritize the mapping locations based on their coverages; or one can lower the parameter for controlling the maximum number of allowed mapping locations for each arm (see details in the Method section) to focus on the uniquely mapped reads. Also, one can annotate the mapping locations using existing genome annotation and identify biological relevant mappings (e.g. in the CLASH study of miRNA-mRNA interactome, the mapping is restricted to the annotated protein-coding genes and miRNA genes [19]).

Existing computational pipelines are available for processing the duplex reads mapping results. For example, Hyb [27] contains scripts for merging the read mappings and detecting genomic islands corresponding to the interacting RNAs, labeling the genomic islands based on existing genome annotation, and performing thermodynamic stability analysis of the predicted RNA duplex *etc*. All the analyses will provide valuable information to assess the biological relevance of the mappings. Most of the existing postprocessing pipelines require BLAST output format or the SAM format [51] as input. Currently, the CLAN output contains information including read ID and RNA arm locations, reference genome chromosome, strand, and exact locations, as well as mapping length *etc.,* which is sufficient to be reformatted into the BLAST output format or the SAM format. Therefore, it is straightforward to couple CLAN with the existing post-processing pipelines to complete the entire analysis. In the near future, we also plan to provide different output formats as options in the new releases of CLAN to allow easier analysis integration and pipeline coupling.

We applied CLAN to re-analyze a public CLASH dataset SRR959751 and identified 40 (out of 46 predicted by CLAN in total) novel miR-10a related interactions, and these novel pathways are involved in pathways relating to cell proliferation regulation and apoptosis with statistical significance (<6*10^−6^). This finding suggests that existing analysis of CLASH data overlooks a significant amount of true interactions due to low mapping rate, and the missed interactions can be retrieved using CLAN, our mapping tool with much higher mapping power. We also note that the miR-10a targets identified here are a subset of all known miR-10a targets, because the CLASH data was generated from a specific cell line. Generating more CLASH data from different cell lines or tissues will reveal a more complete picture of the miRNA interactome.

## Conclusions

In this article, we present a novel duplex read-mapping algorithm CLAN, targeted for analyzing crosslinked RNA sequencing data. To account for the “linker-RNA arm-spacer-RNA arm-linker” pattern of a duplex read, CLAN reformulates a novel computational problem as finding two non-overlapping mappings of the read whose total length is maximized. CLAN exhaustively searches all possible maximal contiguous mappings of any of its prefixes against the reference genome. Then, CLAN merges the mapping according to the reference genome locations to rescue broken mappings due to errors/polymorphisms. Finally, CLAN adopts a dynamic programming-based chaining algorithm to select the two non-overlapping mappings whose total length is maximized. By using BWT and FM-index, CLAN can easily handle the current NGS data volume.

Performance benchmark of CLAN was conducted with BLASTN and STAR on four different crosslinked RNA sequencing technologies, including CLASH, hiCLIP, PARIS, and LIGR-Seq. CLAN was shown to identify much more reads than BLASTN and STAR. In addition to the high mapping sensitivity, the read mapping accuracy of CLAN also appears to match that of BLASTN and higher than that of STAR. In conclusion, the high computational efficiency, high mapping sensitivity, and high mapping accuracy of CLAN make it a powerful tool for crosslinked RNA sequencing data analysis. CLAN is implemented in C++ and freely available on http://sourceforge.net/projects/clan-mapping.

## Methods

### The CLAN algorithm

We formulate the duplex read mapping problem as finding two non-overlapping substrings (i.e. the two RNA arms) whose total mapping length is maximized (see Figure 8 for the high-level summary of the CLAN algorithm).

**Figure 8:**
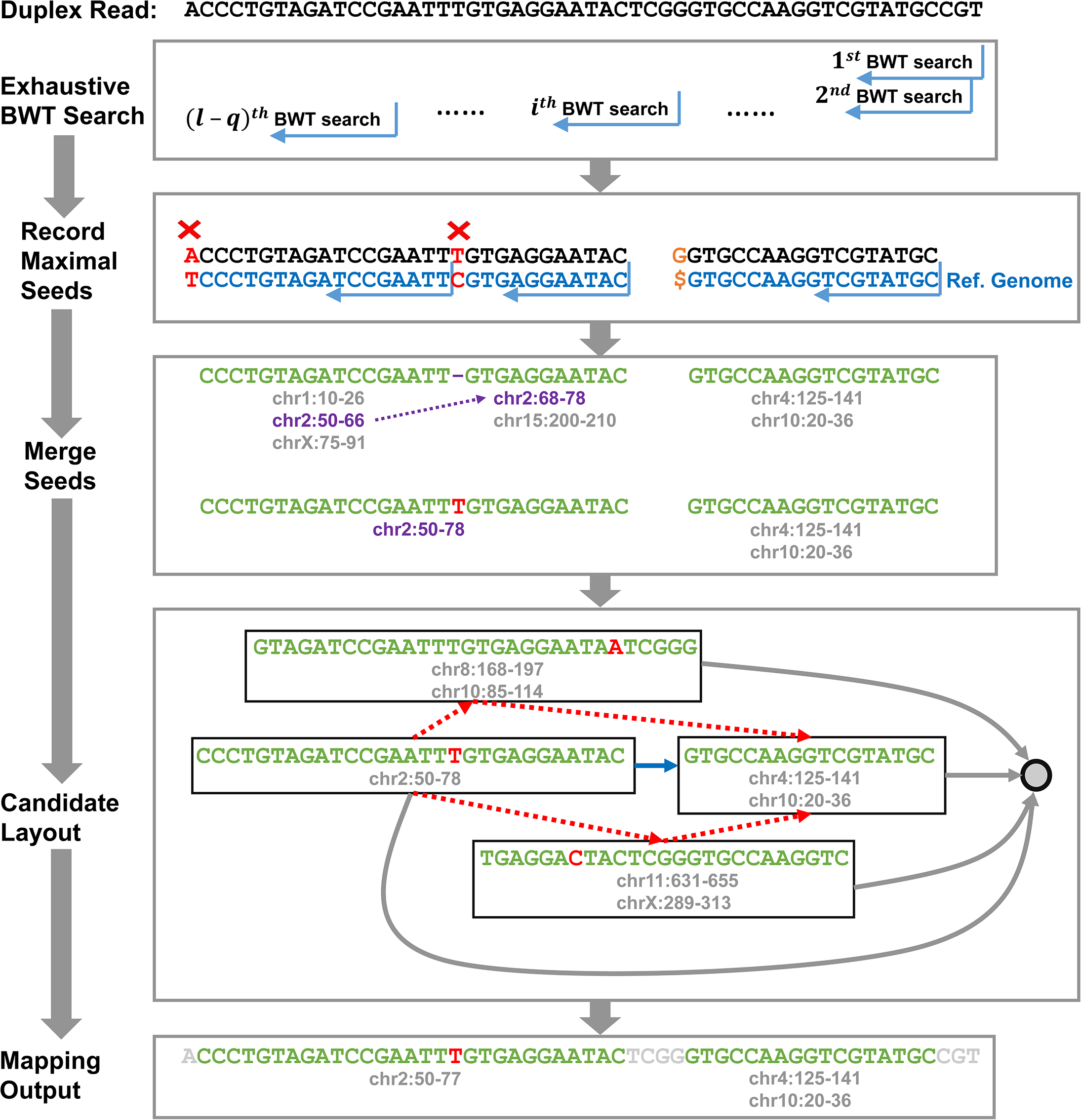
An overview of the CLAN algorithm with an artificial example. The 1^st^ panel “Exhaustive BWT search”: all prefixes of the read are subject to backward BWT search against the constructed reference genome index. The 2^nd^ panel “Record Maximal Seeds”: the BWT search will terminate when a mismatch is encountered (red bases and crosses) or the terminals (an orange base and a terminating symbol “$”) of the indexed strings is reached. The blue sequences correspond to sequences in the reference genome. The 3^rd^ panel “Merge Seeds”: the identified perfect matchings are considered as candidate arms (green strings); each candidate arm is associated with a set of identified reference genome locations. By examining the locations of the candidate arms in the duplex read and their corresponding mapping locations in the reference genome (purple arrow and genome locations), CLAN identifies candidate arm pairs that are potentially broken due to a sequencing error or polymorphism (first row). CLAN merges the candidate arm pairs into a single candidate arm and updates its corresponding mapping location (second row). The 4^th^ panel “Candidate Layout”: a directed acyclic graph (DAG) is generated to represent the relationship between the candidates. Each candidate corresponds to a node (black boxes). Red broken arrows indicate incompatible directed edges due to the overlap between the corresponding nodes; blue solid arrows indicate compatible directed edges; and gray solid edges indicate the directed dummy edges that are added between every node and the dummy terminal (the rightmost gray node). The length of each edge is determined by its source node; and the optimal mapping corresponds to the longest path in the graph that involves no more than two edges. The 5^th^ panel “Mapping Output”: a demonstration of the CLAN output, which contains the two selected candidates and their corresponding genomic locations.

We start the process by first identifying a set of *seeds;* each seed satisfies the following conditions: (1) each seed must be at least *r*nt long (default 10); (2) each seed should be mapped to less than *m* genomics locations (default 20); (3) each seed must be mapped to the reference genome perfectly (no mismatch/gap). CLAN first constructs the BWT and FM-index from the reference genome (as a one-pass step). Then, for a read *s* with length *I,* CLAN performs exhaustive backward BWT search to find all seeds within *s* (see Figure 8, the 1^st^ panel “Exhaustive BWT Search”). For every *r* ≤ *j* ≤ *l* (where *r* is the minimum seed length), CLAN looks for the minimum index *i^j^* such that the substring *s*(*i^j^*,*j*) is a seed. The termination of the extension of a seed could due to an error/polymorphism, or the reach of the termini of the indexed references (see Figure 8, the 2^nd^ panel “Record Maximal Seeds”). To reduce redundancy, the mapping to each genomic location is tracked. A genomic location is considered as *covered* if there exists a backward BWT search that ends on it (or, it is the starting location of a mapping). For each identified seed, all covered genomic positions are subsequently removed from its list of mapped locations. If the list of mapping locations becomes empty, then the entire seed is discarded, otherwise we record the seed and the corresponding mapping locations. Since the direction of the BWT searches is backward, each genomic location will first be covered by the longest seed that begins at this location; only the seeds that are completely contained in the other seeds are eliminated.

The second step is to merge seeds that are potentially broken due to errors/ polymorphisms (see Figure 8, the 3^rd^ panel “Merge Seeds”). For example, in Figure 8, the first two seeds (Figure 8, the 3^rd^ panel, green sequences) map to adjacent locations in the genome (Figure 8, the 3^rd^ panel, first row, purple broken arrow and genomic locations), suggesting that the seeds are potentially broken due to an error/polymorphism. We assume that each RNA arm can be broken for no more than *k* times (by default 1). To describe the merging step, let an arbitrary seed *s*(*i,j*) be mapped to a set of genomic locations, with the *x*th denoted as *T*(*w^x^*,*z^x^*). Here, *T* is the reference genome, and *w^x^* and *z^x^* are the start and end of the mapped genomic interval. For two non-overlapping seeds and *s*(*i*_1_,*j*_1_) and *s*(*i*_2_,*j*_2_) (without loss of generality we assume *i*_2_ > *j*_1_), we attempt to merge the seeds by looking for two adjacent mapped locations, i.e. 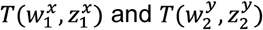 such that:

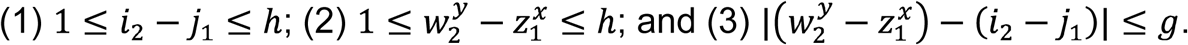

The first two conditions ensure that the two seeds are adjacent in the duplex read and in the reference genome (at most *h*nt apart, default 5); the third condition ensures that the gap (if any) for the corresponding alignment is small (default value of *g* is set to 5). CLAN will exhaustively test all combinations of mapped genomic locations, and merges both seeds into a *candidate* (i.e., *s*(*i*_1_,*j*_2_) with a new mapping location 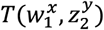, see Figure 8, 3^rd^ panel, second row, purple genomic location) if all conditions are satisfied. A candidate is defined as a substring mapping that contain up to *k* errors/polymorphisms. A seed is by definition a candidate; therefore the candidate set contains all seeds and merged candidates. CLAN iterates this merging process for *k* times, to allow each candidate containing up to *k* errors/polymorphisms.

The third step is to find *f* non-overlapping arms with maximized total mapping length (Figure 8, the 4^th^ panel, “Candidate Layout”). Note that *f* is set to 2 in CLAN for aligning duplex reads; but our algorithm can be extended for any value of *f*. Conceptually, the candidates and their relationships can be represented by a directed acyclic graph (DAG). In the graph, each node corresponds to a candidate (Figure 8, the 4^th^ panel, black boxes). Partially order the candidates based on the increasing order of their starting locations, and break ties with the decreasing order of their ending locations; also consider two nodes as *compatible* if their corresponding candidates do not overlap. For two arbitrary nodes *u* and *v,* a *{u,v}* edge (Figure 8, the 4^th^ panel, blue solid arrow) is added if the following three conditions are satisfied: (1) *u* is partially ordered before *v*; (2) *u* and *v* do not overlap; and (3) no node exists between *u* and *v,* and is simultaneously compatible with both of *u* and *v*. Finally, a dummy node *d* succeeding every other node in the graph (Figure 8, the 4^th^ panel, the rightmost round node) is included into the DAG; and dummy edges are added correspondingly from each node to the dummy node (Figure 8, the 4^th^ panel, gray solid edges). For each edge *{u,v}* CLAN sets its length *l*_*{u,v}*_ as the follows: *l*_*{u,v}*_ = *l_u_* — *c* if *v* ≠ *d* (regular edges); and *l*_*{u,v}*_ = *l_u_* if *v* == *d* (dummy edges). The parameter *c* is the penalty (default 5) for including an additional candidate in the solution set; including this parameter makes the algorithm prefer a single-arm configuration of the read and become more conservative in duplex detection. In this case, the problem of finding two nonoverlapping candidates whose total length is maximized can be transformed as finding the longest path in the DAG that involves no more than *f* edges.

CLAN solves this problem using a dynamic programming (DP) approach. Denote the resulting DAG as *G* = (*V,E*), where *V* corresponds to the node set and *E* corresponds to the edge set. Also let *L*[*v,f*] be the maximum length of the paths which end at *v* and involve *f* edges. CLAN computes *L*[*v,f*] as the follows:

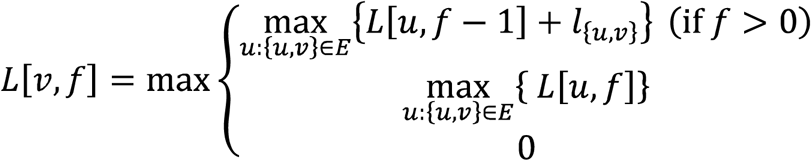

The first condition considers cases where the path is extended to *v* from *u* with the candidate *u* being taken into the solution. The second condition considers similar cases but assumes that the candidate *u* is not taken into the solution. The third condition corresponds to boundary cases where *v* is the starting node of the path. The final solution can be found in *L*[*d,f*], where *d* is the dummy node. The output of the mapping contains the selected arms and their corresponding locations in the duplex read and the reference genomes (Figure 8, the 5^th^ panel, “Mapping Output”).

### Time complexity analysis of the CLAN algorithm

Denote the length of a duplex read as *l*. Clearly, with the help of BWT and FM-index, the search of an *l-*long sequence against the reference genome requires *O*(*l*) time. Because CLAN searches every prefix of the duplex read and there are at most *l* prefixes, the total time required for the exhaustive BWT search step adds up to *O*(*l*^2^). For candidate merging, CLAN tests the merging of every pair of candidate seeds in the worst-case scenario, which leads to an *O*(*m*^2^*l*^2^) complexity, where *m* is the number of genomic locations associated with each candidate. However, because *m* is a constant (by default 20), and CLAN only attempts to merge adjacent candidates (parameter *h*, by default 5), the merging step is in fact very efficient. Finally, for the DP-based chaining, each node *v* has at most *l* nodes that precede it; as a result, computing the answer for each DP-table entry requires *O*(*l*) time. There are *O*(*fl*) entries of the DP table *L,* and the total time required for the chaining step is thus *O*(*fl*^2^). Since *f* is set to 2 for duplex read mapping, the time complexity for the DP chaining step is *O*(*l*^2^). Taken together, CLAN requires *O*(*l*^2^) to map a single duplex read. Note that the duplex read length *l* is technology-dependent and can also be considered as a constant; CLAN thus requires a constant time to map a single duplex read, and the overall running time is linear with respect to the throughput of the experiment (or the number of reads in the dataset).

### Computing Interests

The authors declare that they have no competing interests.

